# Large oligomeric STOML3 rings scaffold mechanosensitive ion channels

**DOI:** 10.64898/2026.11.23.768286

**Authors:** Angela Tzu-Lun Huang, Clement Verkest, James Hall, Max Ruwolt, Fan Liu, Stefan G. Lechner, Gary R. Lewin

## Abstract

Stomatin-like protein 3 (STOML3) is a protein required for touch transduction and is a member of the SPFH (Stomatin/Prohibitin/Flotillin/HflKC) superfamily. STOML3 strongly regulates the force sensitivity of both PIEZO and ELKIN1 ion channels, yet its molecular mode of action is not understood. Multi-scale super-resolution imaging revealed that STOML3 assembles into very large heterogeneous ring-like structures (0.1–1.6 µm diameter) at cholesterol-enriched membrane domains consisting of protomers with a Type II transmembrane topology. Ring formation depends on a conserved proline (P44), and its mutation (P44S) abolished ring architecture without eliminating membrane targeting. STOML3 physically associates with PIEZO1, PIEZO2, and ELKIN1 at these domains and restricts the mobility of PIEZO2 clusters. Disrupting ring formation by mutation, by pharmacological inhibition of oligomerization, or by cholesterol depletion reduced channel association. These findings identify STOML3 rings as large supramolecular assemblies that organize mechanosensitive ion channels at the plasma membrane.

## Introduction

Rapid detection of extremely small mechanical forces is a fundamental property of sensory systems in all animals^1^. This process, called fast mechanotransduction, relies on the gating of ion channels by mechanical force. Interestingly, the ion channels that are necessary for sensory transduction are molecularly diverse in worms, flies and mammals^2–6^. However, one unifying theme across these sensory systems is that accessory proteins have been identified and are often necessary to tune the sensitivity of the transduction process. Such proteins were first identified in the nematode *Caenorhabditis elegans* (*C. elegans*) and may be membrane proteins that interact with the pore-forming ion channel, cytoskeletal elements, or proteins that tether mechanosensitive ion channels to the cytoskeleton or extracellular matrix^7–10^. The body touch receptor of the nematode is probably the best-characterized mechanoreceptor neuron, as genetic screens have identified essentially all proteins necessary for light touch transduction on the body^11,12^. The pore-forming mechanosensitive ion channel is composed of MEC-4 and MEC-10, both members of the DEG/ENaC superfamily^2,13^. Mutations that change the ion selectivity of MEC-4 channels also change the biophysical properties of mechanosensitive currents in vivo^2,14^, providing strong evidence that these channels form the ion conduction pathway for touch transduction. Although as many as 17 proteins may be involved in the physiological assembly of the transduction apparatus^12,13^, only a handful are absolutely necessary for transduction currents. These include the pore-forming channel subunit MEC-4, but also the stomatin-domain protein MEC-2^2^. MEC-2 belongs to the large, evolutionarily conserved SPFH (Stomatin, Prohibitin, Flotillin, HflK/C) superfamily^15–17^, whose members have highly diverse roles across all kingdoms of life^18^. In *C. elegans*, MEC-2 physically interacts with the DEG/ENaC channels MEC-4 and MEC-10 to regulate channel activity^2,9,14^.

Two closely related mammalian orthologues of MEC-2 are stomatin and stomatin-like protein-3 (STOML3)^18^. Stomatin was first identified as a protein missing from the plasma membrane of human erythrocytes from patients with stomatocytosis^19–23^. However, mice lacking either stomatin or STOML3 show deficits in sensory mechanotransduction indicating that these proteins may have a conserved role in regulating mechanotransduction across evolution^20,24^. However, while genetic loss of *stomatin* has a relatively small effect on cutaneous mechanoreceptor sensitivity in the mouse^20^, loss of STOML3 leads to the complete loss of mechanotransduction in almost half of all cutaneous mechanoreceptors^24–26^. We determined that loss of sensory mechanotransduction in *Stoml3^-/-^* mice is likely due to the fact that STOML3 can lower the threshold for activation of mechanosensitive ion channels required for touch, like PIEZO2 and ELKIN1^6,27–29^. Indeed, the deficits in mechanoreceptor function seen in *Piezo2*, *Elkin1* and *Stoml3* mutant mice are very similar to each other, indicating that these proteins cooperate to regulate sensory mechanotransduction^5,6,25,26,30^. Nevertheless, the mechanism by which STOML3 lowers the amount of force needed to activate mechanosensitive channels remains unclear. STOML3 molecules are associated with cholesterol rich lipid domains and may thereby mediate local membrane stiffening^25,31,32^. There is also good evidence that STOML3 forms oligomers although the precise structural basis of such complexes has remained unclear^18,25,32,33^. Importantly, mutations in the STOML3 protein that disrupt STOML3 self-association or small molecules that inhibit the proteins oligomerization are sufficient to abolish the functional modulation of ion channels both *in vivo* and *in vitro*^25,33^. In contrast to the ubiquitously expressed *stomatin*^34^, *stoml3* shows a relatively restricted expression pattern with expression observed in dorsal root ganglia^24,25^, but even higher levels observed in olfactory sensory neurons^24,25,35,36^. However, deficits in olfactory sensory neuron function are mild in *Stoml3* mutant mice^35,37^.

Recent cryo-EM structures of a variety of SPFH family members has revealed a common architectural feature of this family, namely that they form ring or dome-like structures in the membrane^16,17,38–43^. In the case of human stomatin, so far three cryo-EM structures show an oligomeric dome like structure consisting of 16 monomers^39,41,43^. The dome-like stomatin structure has an internal diameter of ∼120 Å and an external width of ∼160 Å, but a much narrower opening to the dome on the intracellular side^39,41^. Structures determined from heterologously expressed protein are also almost identical to the domes purified from native red blood cell membranes^39,41,43^. Initial biochemical characterization of the stomatin protein indicated that both the N- and C-terminus of the protein are intracellular which is consistent with recent structures^43,44^. However, it has also been suggested that alternative single-pass forms of stomatin exist^44^. These studies naturally raise the question whether the STOML3 protein forms a similar dome-like structure in the membrane and whether ring-like structures of STOML3 could enclose ion channels like stomatin does in the red blood cell membrane^17,41^.

Here, we determined the membrane topology of STOML3 and its organization in the membrane using super-resolution microscopy in cell lines and primary sensory neurons. We show that a conserved proline residue is required for higher-order ring-like cluster formation. STOML3-rings appear to have a membrane topology distinct from stomatin and can form ring structures an order of magnitude larger than those of stomatin and other SPFH proteins. We show that STOML3 rings concentrate PIEZO1, PIEZO2 and ELKIN1 at the plasma membrane and restrict the lateral mobility of PIEZO2 clusters. Finally, we show that manipulations that disrupt ring assembly, reduce the association of STOML3 with these channels, providing a mechanism for the modulation of sensory mechanotransduction.

## Results

### STOML3 is a lipid raft protein with Type II transmembrane topology

We first studied STOML3 membrane organization by examining its association with lipid raft microdomains using sucrose density gradient fractionation. We generated a C-terminal ALFA-tagged cDNA (STOML3-ALFA) which we expressed in HEK293T cells. Overexpressed STOML3 co-fractionated with endogenous Flotillin-2, a well-established lipid raft marker^46,47^, in low-density fractions (Fig. 1a), confirming that STOML3 preferentially localizes to cholesterol-enriched membrane domains.

**Figure 1.**
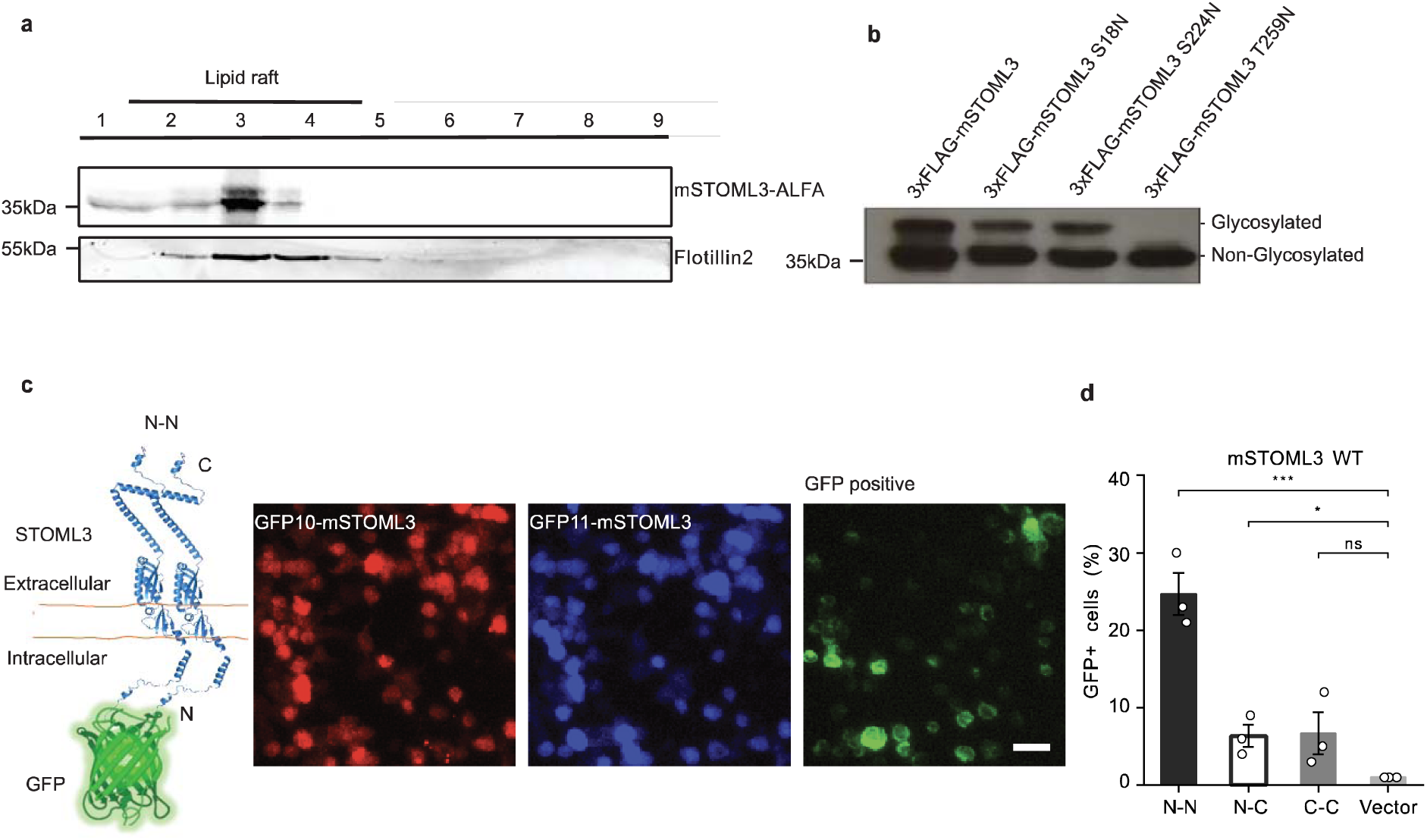
STOML3 localizes to lipid rafts and adopts a Type II transmembrane topology. **(a)** Sucrose gradient fractionation of HEK293T cells expressing mSTOML3-ALFA. STOML3 co-fractionates with the lipid raft marker Flotillin-2 in low-density fractions 2-5 (lipid rafts). Molecular weight markers: STOML3-ALFA, 35 kDa; Flotillin-2, 55 kDa. **(b)** N-glycosylation site mapping by western blot. Wild-type mSTOML3 and point mutants (S18N, S224N, T259N) were expressed in HEK293T cells as 3xFLAG-tagged constructs; the plasma membrane fraction was analysed and detected with anti-FLAG (M2)-HRP. The T259N mutation eliminates the glycosylated band, indicating extracellular localization of this region and confirming the C-terminus is extracellular. **(c)** Representative images of the N-N tripartite split-GFP topology assay for wild-type mSTOML3. Schematic (left) illustrates the N-N configuration, with GFP10 and GFP11 fragments fused to the N-termini of two STOML3 molecules and the GFP1-9 detector expressed in the cytoplasm. GFP complementation occurs when both N-termini reside in the cytoplasm. Microscopy images show GFP10-mSTOML3 (red), GFP11-mSTOML3 (blue), and resulting GFP complementation signal (green). Scale bar: 20 μm. **(d)** Quantification of tripartite split-GFP topology assay for wild-type mSTOML3. GFP-positive cells were quantified for N-N, N-C, C-C, and vector control combinations. High GFP signal in N-N pairs and low signal in N-C and C-C pairs confirms Type II topology (N-terminus cytoplasmic, C-terminus extracellular). Data are mean ± SEM from n = 3 independent experiments. ***P < 0.001, *P < 0.05; n.s., not significant (one-way ANOVA with Tukey’s post-hoc test).

Previous studies demonstrated that stomatin-domain family members may exist in two topological states: a hairpin-loop form with both termini cytoplasmic, and an N-glycosylated single-pass transmembrane form^44^. In stomatin, the hairpin-loop form predominates, with the transmembrane form representing a minor population^44^. Recent cryo-EM structures of stomatin captured this major hairpin-loop population in an oligomeric dome^39,41,43^. To establish the membrane topology of the highly related STOML3, we used two complementary approaches: N-glycosylation site mapping and a tripartite split-GFP complementation assay. The consensus sequence for N-glycosylation (NXS/T, where X is any amino acid except proline)^48,49^ occurs at three positions in the mouse STOML3 sequence: residues 16-18, 222-224, and 257-259. We introduced point mutations (S18N, S224N, and T259N) to abolish each potential glycosylation site and assessed their glycosylation status by Western blot. Wild-type STOML3 appeared as two bands: a higher molecular weight glycosylated form and a lower molecular weight form (Fig. 1b). To confirm that the upper band reflects N-glycosylation rather than another post-translational modification, lysates were split and treated with PNGase F, which collapsed the doublet to the lower band (Supplementary Fig. 1b), as previously described for stomatin^44^. The T259N mutation, which removes the C-terminal N-glycosylation consensus site, eliminated the upper band, confirming that this site is glycosylated and therefore exposed to the ER lumen/extracellular space. The persistence of the lower band in wild-type STOML3 could reflect either a minor hairpin-loop population (where the C-terminus remains cytoplasmic and cannot be glycosylated) or incomplete glycosylation of the Type II form. Western blots of endogenous STOML3 from sciatic nerve of StrepII-STOML3 knock-in mice also showed the same higher molecular weight bands indicating that the endogenous protein is likely glycosylated^25^. Sequence alignment revealed that this glycosylation site is conserved across STOML3 orthologs (T259 in mouse, T263 in human; Supplementary Fig. 1e).

To directly determine membrane topology, we employed a tripartite split-GFP complementation assay in which GFP fragments (GFP10 and GFP11) are fused to the N- or C-terminus of STOML3, with a cytoplasmic GFP1-9 detector (Supplementary Fig. 1a). GFP complementation occurs only when both tagged termini are in the cytoplasm (Fig. 1c). We observed a robust GFP signal for N-N STOML3 pairs (∼25% GFP-positive cells) significantly higher than vector controls but a minimal signal for N-C and C-C combinations (Fig. 1c, d). This pattern is uniquely consistent with Type II transmembrane topology, where the N-terminus faces the cytoplasm and C-terminus is extracellular (Fig. 1d; Supplementary Fig. 1c). Importantly, the T259N glycosylation mutant showed an identical complementation pattern to wild-type STOML3 (Supplementary Fig. 1d), confirming that this mutation does not alter membrane topology.

Together, these data establish that STOML3 predominantly adopts a Type II transmembrane topology, preferentially localizing to lipid raft membrane domains. While we cannot exclude a minor hairpin-loop population, the robust extracellular ALFA-tag detection (Fig. 2) and split-GFP complementation pattern indicate that the Type II form represents the major STOML3 population in contrast to stomatin, where the hairpin-loop form predominates^39,44^.

**Figure 2.**
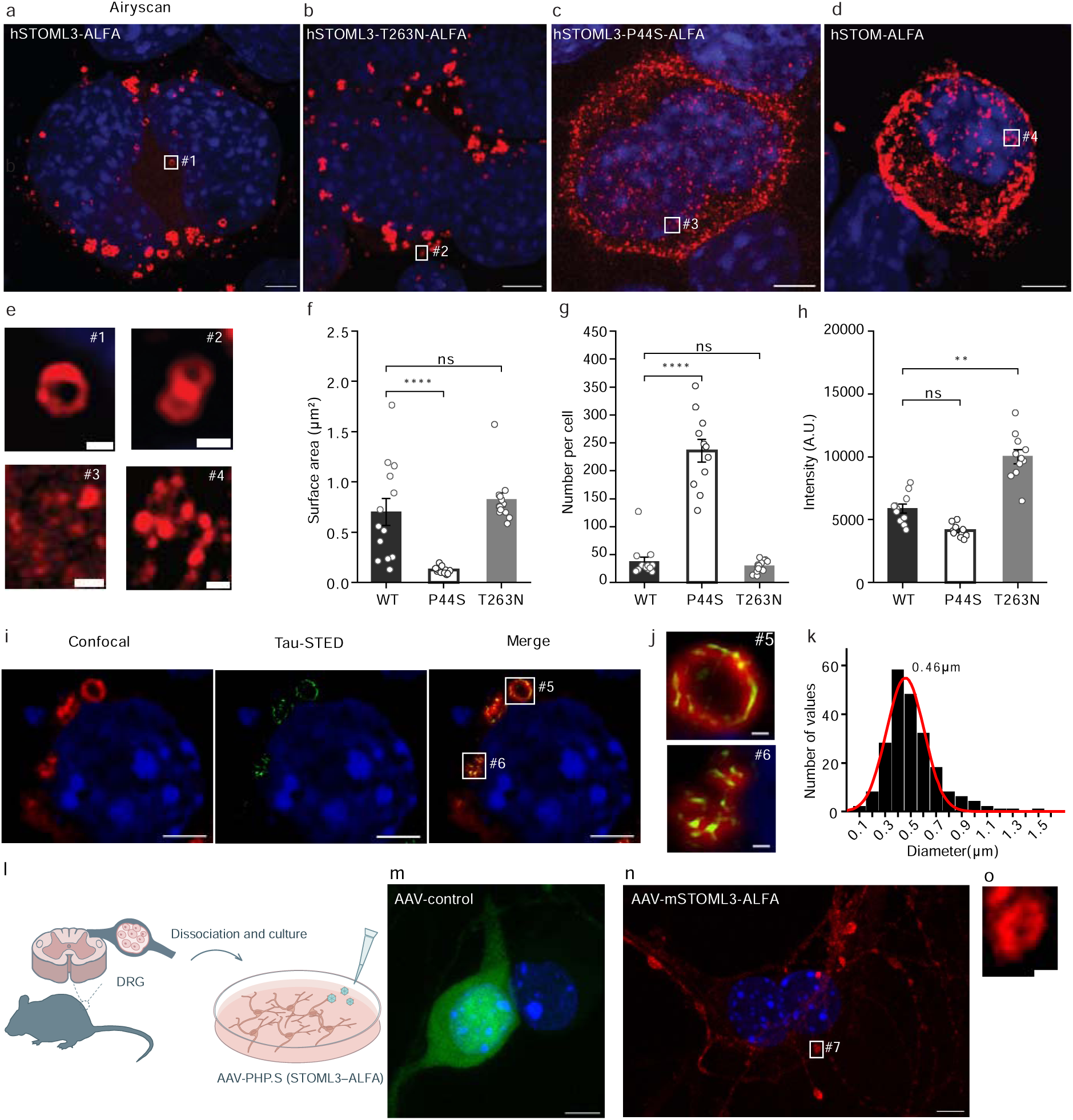
STOML3 forms ring-like membrane structures dependent on a conserved membrane-proximal proline. **(a-d)** Airyscan microscopy of ALFA-tagged proteins in non-permeabilized N2a cells, detecting extracellular epitopes. **(a)** hSTOML3-ALFA forms distinctive ring-like clusters at the plasma membrane. **(b)** T263N glycosylation mutant retains ring formation. **(c)** P44S mutation in the membrane-proximal cholesterol-binding region completely disrupts ring formation, showing scattered membrane distribution. **(d)** Human stomatin (hSTOM-ALFA) does not form ring-like structures under identical conditions. **(e)** Magnified insets (#1-4) showing ring morphology for each construct. **(f-h)** Quantification of cluster parameters (n = 10 cells per condition from 3 independent experiments): surface area **(f)**, cluster number per cell **(g)**, and fluorescent intensity **(h)**. P44S shows significantly reduced surface area (****P < 0.0001) and cluster number (****P < 0.0001) compared to WT; T263N shows comparable surface area and cluster numbers but elevated intensity (**P < 0.01). **(i-k)** Tau-STED microscopy confirms ring structures with improved resolution. **(i)** Confocal (left), Tau-STED (center), and merged (right) images of hSTOML3-ALFA at the plasma membrane. DAPI nuclear staining in blue. **(j)** Magnified insets (#5, #6) showing individual ring morphology. **(k)** Distribution of ring diameters (n = 255 rings from 9 cells, 3 independent experiments), ranging from 0.1-1.6 μm with a peak at 0.46 μm. **(l)** Schematic of AAV-PHP.S-mediated expression of mSTOML3-ALFA in dissociated mouse DRG neuron cultures. **(m-o)** Airyscan images of non-permeabilized DRG neurons transduced with AAV-control **(m)** or AAV-mSTOML3-ALFA **(n)**, with magnified inset (#7) showing ring-like clusters at the neuronal surface **(o)**. Scale bars: 5 μm (overview), 500 nm (insets). Statistical comparison by one-way ANOVA with Dunnett’s post-hoc test. Images representative of 3 independent cultures.

### STOML3 forms very large oligomeric ring-like membrane structures

Recent cryo-EM structures have revealed that SPFH family members such as stomatin and flotillin assemble into closed, uniform oligomeric cages consisting of 16 (stomatin) to 44 (flotillin) subunits^16,17,38–41,43^, yet it was unknown whether STOML3 adopts a similar architecture. To visualize the membrane organization of STOML3 by super-resolution microscopy, we developed an imaging strategy using the STOML3-ALFA construct, enabling detection with nanobody-conjugated fluorophores in non-permeabilized cells. This approach provides independent validation of the extracellular C-terminus established from our topology studies (Fig. 1). Using Airyscan microscopy (lateral resolution ∼120 nm), we observed distinctive ring-like formations of STOML3 at the plasma membrane (Fig. 2a, e), structures considerably larger than the closed cages reported for human stomatin^39,41,43^. Quantitative analysis of more than 10 randomly selected cells from three independent experiments revealed that these ring-like clusters exhibited a mean surface area of approximately 0.7 μm² and occurred at a frequency of ∼30 clusters per cell (Fig. 2f, g). Notably, Airyscan imaging revealed considerable heterogeneity in cluster dimensions, suggesting that STOML3 rings may not be a single uniform species. Mouse STOML3 showed similar ring formation to human STOML3, with comparable cluster surface area and numbers (Supplementary Fig. 2e-g). Almost identical ring-like structures were also observed in the T263N glycosylation mutant, with cluster surface area and numbers comparable to wild-type (Fig. 2b, f, g). However, fluorescent intensity was significantly elevated in T263N-expressing cells (Fig. 2h), suggesting that N-glycosylation may modulate STOML3 cluster stability or membrane retention without affecting overall ring architecture.

A conserved proline residue in the membrane proximal region of MEC-2, mouse STOML3, and human STOML3 has been implicated in cholesterol binding and is critical for protein function (P134 in MEC-2, P40 in mouse, P44 in human; Supplementary Fig. 2a)^31,50,51^. In stomatin, the equivalent proline is essential for hairpin-loop topology and cholesterol binding^44,45^, and in mouse STOML3, the P40S mutation abolishes lipid raft localization, eliminates the ability to potentiate PIEZO channel currents, and prevents rescue of mechanosensitivity in STOML3-deficient neurons^31^. We therefore hypothesized that this residue would also be critical for the ring-like supramolecular organization of STOML3.

Indeed, the P44S mutation completely disrupted large ring formation, with STOML3 appearing scattered across the membrane rather than in organized clusters (Fig. 2c, e). Quantification revealed that while membrane targeting was preserved, STOML3 was redistributed from large ring-like clusters to scattered puncta: individual cluster surface area decreased dramatically (approximately 0.11 μm² compared to 0.72μm² for wild-type) accompanied by a striking ∼7-fold increase in the number of dispersed puncta (approximately 224 per cell compared to 31 for wild-type) (Fig. 2f, g), indicating that the membrane targeting is not abolished, but redistributed, and the cholesterol-binding proline is essential for the supramolecular assembly of STOML3 into ring structures. These effects were also seen for the mouse STOML3, where the P40S mutation similarly disrupted ring formation (Supplementary Fig. 2b-d). Notably, the closely related SPFH family member stomatin did not form large ring-like structures under identical imaging conditions (Fig. 2d, e), demonstrating that this organizational property is specific to STOML3 rather than a general property of proteins with an SPFH domain.

To validate these observations at higher resolution and further characterize the size and heterogeneity of STOML3-rings, we employed STED (Stimulated Emission Depletion) microscopy (lateral resolution ∼40–60 nm). Tau-STED imaging, which combines stimulated emission depletion with fluorescence lifetime gating for improved signal-to-noise ratio, confirmed the presence of discrete ring structures consistent with Airyscan results (Fig. 2i, j). Because the axial resolution of both Airyscan and STED is several hundred nanometres, these images represent projections of STOML3 structures along the optical axis. Quantitative analysis of ring diameters by STED microscopy on 255 rings from 9 cells across three independent experiments revealed a broad size distribution ranging from 0.1 to 1.6 μm, with a peak diameter of 0.46 μm (Fig. 2k), again suggesting that STOML3 assembles into rings of variable dimensions. Furthermore, expression of mouse STOML3 in both N2a wild-type and STOML3KO cell lines produced indistinguishable ring clusters (Supplementary Fig. 2h-j)^52^, confirming that ring formation is an intrinsic property of STOML3 independent of endogenous protein levels.

To examine whether STOML3 forms comparable structures in sensory neurons, we transduced cultured mouse dorsal root ganglion (DRG) neurons with AAV-PHP.S encoding mSTOML3-ALFA (Fig. 2l)^6^. Surface labelling of non-permeabilized neurons revealed ring-like STOML3 clusters at the neuronal plasma membrane, whereas no ALFA signal was detected in neurons transduced with control AAV (Fig. 2m-o). To exclude the possibility that these clusters arise during fixation, we labelled the ALFA-tag in living cells before fixation. Under these conditions, hSTOML3-ALFA formed ring-like clusters in N2a cells, whereas hSTOML3-P44S-ALFA remained scattered across the membrane. Large ring-like clusters were also observed in live-labelled DRG neurons (Supplementary Fig. 3). Because the nanobody was applied to intact living cells, these data also independently support an extracellular STOML3 C-terminus. Together, these results indicate that ring-like cluster formation is neither restricted to heterologous cell lines nor a consequence of fixation.

### MINFLUX nanoscopy resolves STOML3 rings at molecular resolution

To examine STOML3 rings at molecular resolution, we combined 3D-MINFLUX nanoscopy with DNA-PAINT, using an anti-GFP nanobody carrying a DNA docking strand to label N-terminally EGFP-tagged hSTOML3 expressed in N2a cells (Fig. 3b). Because the EGFP tag is cytoplasmic, cells were permeabilized for labelling. Localization precision was approximately 3 nm in all three dimensions (Fig. 3d).

**Figure 3.**
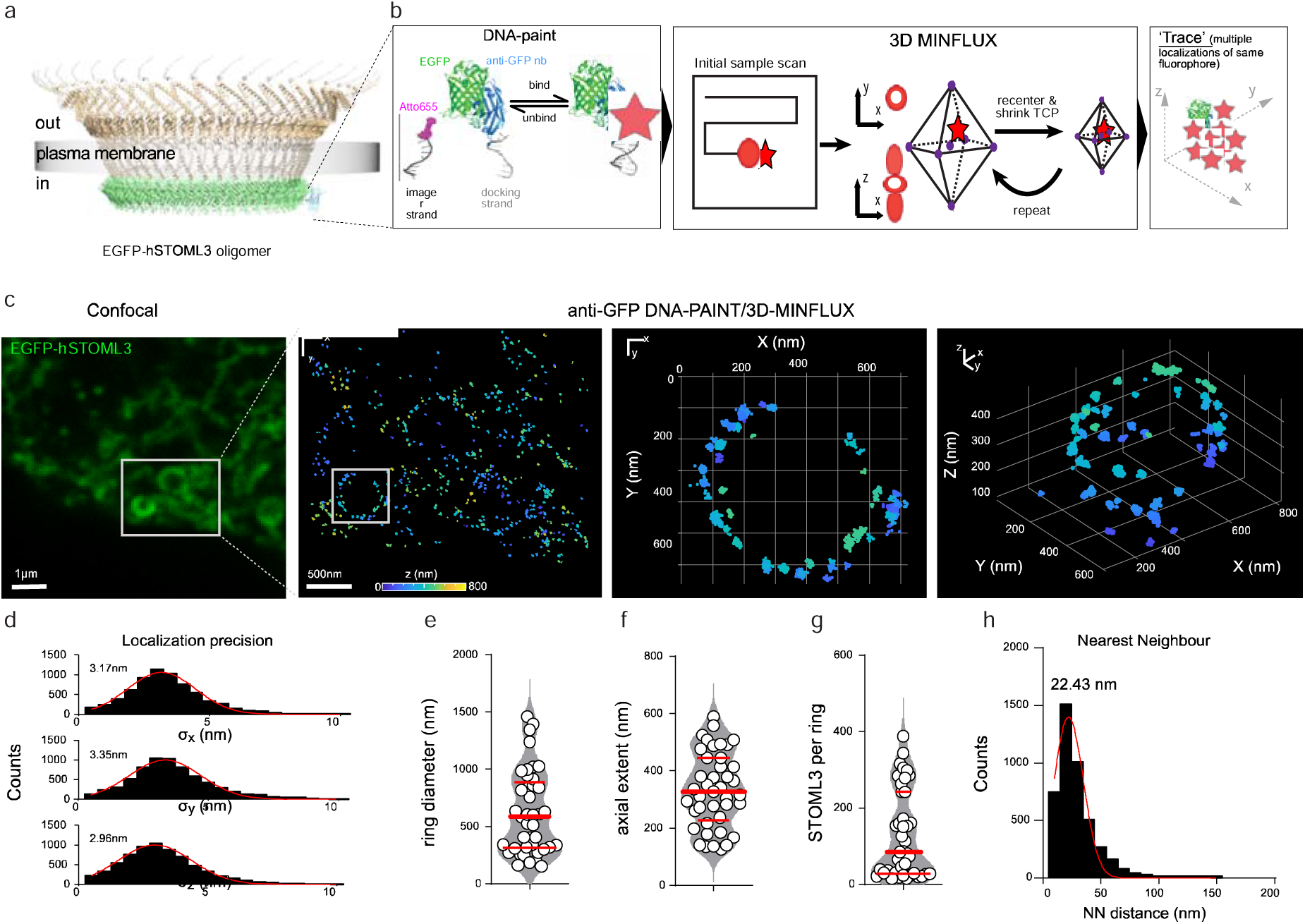
MINFLUX nanoscopy resolves STOML3 rings at molecular resolution. **(a)** Model of an EGFP-hSTOML3 oligomer in the plasma membrane, with the N-terminal EGFP tags on the cytoplasmic side. **(b)** Labelling and imaging strategy. An anti-GFP nanobody (nb) carrying a DNA docking strand binds EGFP-hSTOML3; transient hybridization of an Atto655-labelled imager strand enables iterative 3D-MINFLUX localization, in which the targeted coordinate pattern (TCP) is recentered and contracted around the emitter. Repeated localizations of the same fluorophore are grouped into a trace. **(c)** N2a cells expressing EGFP-hSTOML3, fixed and permeabilized. From left: confocal image; MINFLUX image of the boxed region, colour-coded by axial (z) position (0–800 nm); xy projection of the ring boxed in the MINFLUX image; 3D view of the same ring. Scale bars, 1 µm (confocal) and 500 nm (MINFLUX). **(d)** Localization precision in x, y and z; red lines, Gaussian fits (peaks σx = 3.17 nm, σy = 3.35 nm, σz = 2.96 nm; n = 9,226 localizations). **(e–g)** Ring diameter **(e)**, axial extent **(f)** and number of detected STOML3 molecules per ring **(g)**; each circle represents one ring; thick red line, median; thin red lines, interquartile range. **(h)** Nearest-neighbour distances between detected STOML3 molecules within rings (n = 4,726 distances); red line, Gaussian fit (peak 22.43 nm). n = 41 rings from 3 cells in three independent experiments.

MINFLUX resolved EGFP-hSTOML3 as discrete clusters at the plasma membrane– coverslip interface, many of which formed rings resembling those observed by Airyscan and STED microscopy (Fig. 3c). Across 41 rings from three independent experiments, ring diameters ranged from 153 to 1,463 nm (median 587 nm, interquartile range 316– 889 nm; Fig. 3e), comparable to the size distribution measured by STED (Fig. 2k). Rings contained a median of 84 detected STOML3 molecules (range 15–388; Fig. 3g). Because DNA-PAINT labelling and detection are incomplete, these values represent lower estimates of the number of STOML3 molecules per ring. Within rings, detected molecules were closely spaced: the nearest-neighbour distance distribution peaked at approximately 22 nm, and 86% of molecules had a neighbour within 50 nm (Fig. 3h).

The axial extent of the rings (median 328 nm, range 127–589 nm; Fig. 3f) exceeded that expected for a single layer of membrane-bound STOML3. Because labelling of the cytoplasmic EGFP tag required permeabilization, this axial spread may reflect local plasma membrane topography, labelled STOML3 in intracellular membranes close to the cell surface, or both, which cannot be distinguished in these experiments. The surface localization of STOML3 rings is, however, independently established by labelling of the extracellular ALFA tag in non-permeabilized and living cells (Fig. 2; Supplementary Fig. 3). Even the smallest rings resolved by MINFLUX (153 nm in diameter) were approximately ten times wider than the 16-mer stomatin dome (∼16 nm) ^39,41^, indicating that STOML3 rings are far larger than any SPFH assembly characterized structurally so far.

### STOML3 co-fractionates and biochemically interacts with mechanosensitive ion channels

Previous studies have demonstrated that STOML3 functionally modulates mechanosensitive ion channels including PIEZO1, PIEZO2, and ELKIN1^6,27,28,31^. However, the molecular basis for this functional association remained unclear. Having established the membrane topology and supramolecular organization of STOML3, we next investigated its interactions with mechanosensitive ion channels.

Sucrose gradient fractionation revealed that PIEZO2 and ELKIN1 co-fractionated with STOML3 in lipid raft fractions 2–5 (Fig. 4a, Supplementary Fig. 4a), demonstrating that these proteins reside in the same membrane microdomains. To test for a biochemical interaction, we performed affinity purification using STOML3-ALFA. When expressed in HEK293T cells, human STOML3 co-purified with all tested mechanosensitive ion channels (PIEZO1, PIEZO2 and ELKIN1; Fig. 4b). Mouse STOML3 showed identical interaction profiles (Supplementary Fig. 4b). In contrast, stomatin failed to co-purify with either PIEZO1 or PIEZO2 under the same conditions (Supplementary Fig. 4d), indicating that the ability to interact with mechanosensitive channels is specific to STOML3. To identify STOML3 interactors in an unbiased manner, we expressed hSTOML3-ALFA in HEK293T cells, immunoprecipitated it with ALFA-Selector beads, and analyzed the bound proteins by mass spectrometry. Volcano plot analysis identified 1,393 significantly enriched proteins (P < 0.02, |log₂ fold-change| > 1). Endogenous ELKIN1 (TMEM87A) was among the most enriched interactors (Fig. 4e), supporting the STOML3–ELKIN1 interaction at endogenous ELKIN1 expression levels. Gene Ontology analysis of the interactome revealed enrichment for endoplasmic reticulum membrane and mitochondrial components, protein transport and folding, and ribosomal and translation-associated proteins (Supplementary Fig. 4c). It should be noted that such pull down experiments will reveal all possible interactions of STOML3 during its lifetime, including those during its biogenesis, membrane trafficking and at the plasma membrane.

**Figure 4.**
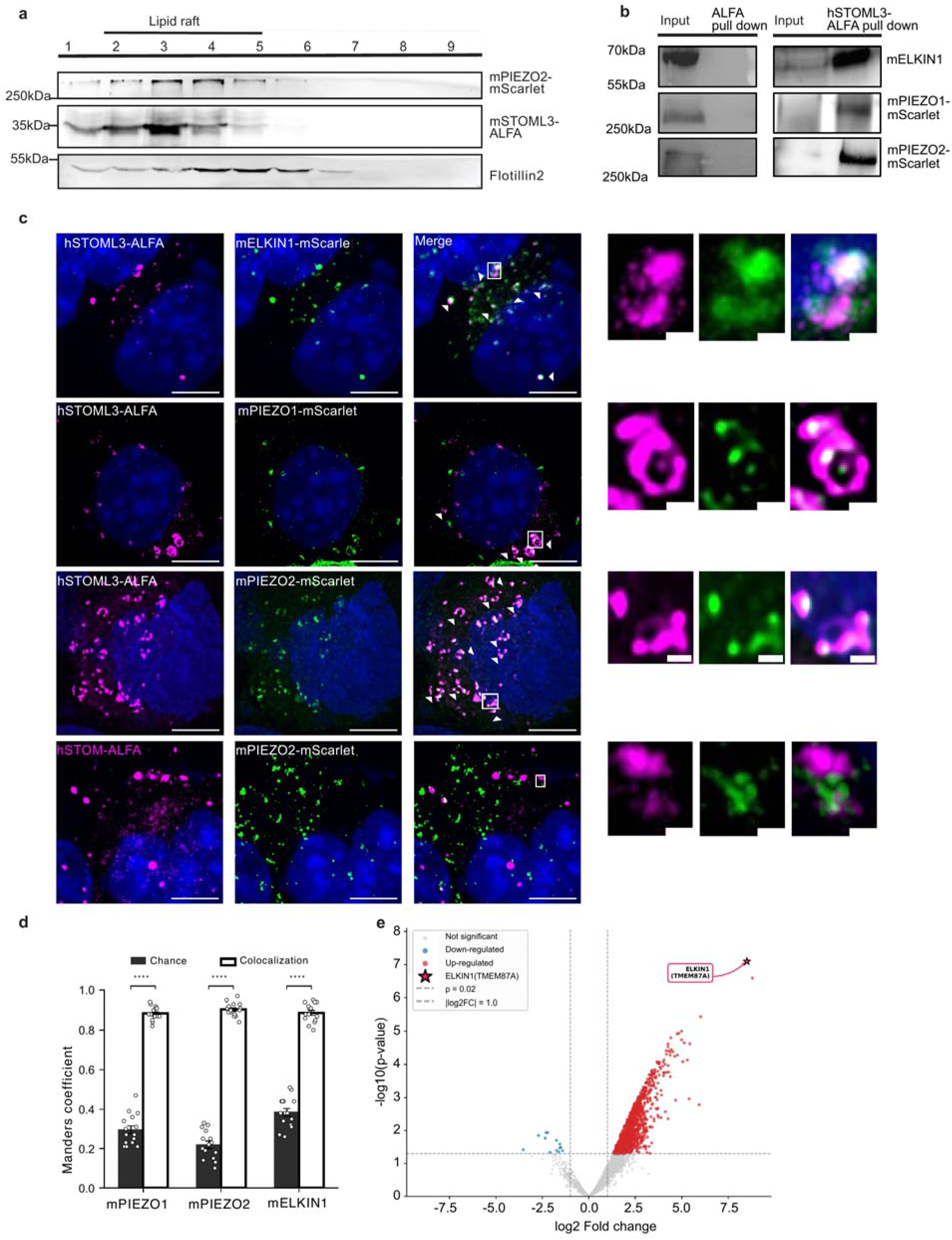
STOML3 co-localizes and physically associates with mechanosensitive ion channels at lipid rafts. **(a)** Sucrose gradient fractionation showing co-fractionation of mSTOML3-ALFA, mPIEZO2-mScarlet, and Flotillin-2 in lipid raft fractions 2-5. Molecular weight markers: STOML3-ALFA, 35 kDa; PIEZO2-mScarlet, 300 kDa; Flotillin-2, 55 kDa. **(b)** Affinity purification demonstrating association between hSTOML3-ALFA and mechanosensitive channels. ALFA-tag pull-down co-purifies mELKIN1-mScarlet (70 kDa), mPIEZO1-mScarlet (300 kDa), and mPIEZO2-mScarlet (300 kDa). Input and pull-down fractions shown. **(c)** Airyscan colocalization of hSTOML3-ALFA (magenta) with mELKIN1-mScarlet, mPIEZO1-mScarlet, and mPIEZO2-mScarlet (green) at the plasma membrane. hSTOM-ALFA (stomatin) with mPIEZO2-mScarlet serves as negative control. DAPI nuclear staining in blue. Merge panels show colocalization; arrowheads indicate co-localizing structures. Right panels show magnified insets of boxed regions with individual channels and merge. Scale bars: 5 μm (overview), 500 nm (insets). **(d)** Quantification of Manders’ colocalization coefficients for STOML3 with PIEZO1, PIEZO2, and ELKIN1. All three channels show significantly higher colocalization with STOML3 compared to chance levels (****P < 0.0001, unpaired t-test). **(e)** Mass spectrometry volcano plot from hSTOML3-ALFA immunoprecipitation in HEK293T cells. Endogenous ELKIN1 (TMEM87A) is identified as one of the most significantly enriched interactors (highlighted). Significance cutoffs: P = 0.02, |log₂ fold-change| = 1.0. Red, up-regulated; blue, down-regulated; gray, not significant. Data are mean ± SEM from n = 3 independent experiments (a-d). MS data from n = 3 biological replicates (e).

Airyscan imaging demonstrated colocalization of mechanosensitive channels with STOML3 ring clusters at the plasma membrane (Fig. 4c). Quantitative analysis using Manders’ coefficients revealed highly specific colocalization (approximately 0.9) between STOML3 and each of the three mechanosensitive channels (PIEZO1, PIEZO2, and ELKIN1), significantly exceeding chance overlap levels of approximately 0.2–0.4 (Fig. 4d). Consistent with the biochemical data, stomatin showed no colocalization with PIEZO2 under identical imaging conditions (Fig. 4c, lower panels), further demonstrating that channel association is a specific property of STOML3 rather than a general feature of SPFH domain proteins.

To examine the dynamic relationship between STOML3 and mechanosensitive channels, we performed live-cell TIRF imaging of PIEZO2-HaloTag with STOML3-GFP. PIEZO2 clusters tracked together with STOML3 clusters at the plasma membrane, demonstrating their co-movement in real time (Supplementary Fig. 5a). To test whether STOML3 constrains the mobility of PIEZO2 clusters, we compared PIEZO2-HaloTag diffusion in N2a-PIEZO1KO cells (with STOML3) and N2a-STOML3KO cells. While cluster diameter, density, and overall track distribution were unchanged (Supplementary Fig. 5b, d), the absence of endogenous STOML3 was associated with increased diffusion coefficients of sub-diffusive and confined PIEZO2 clusters—populations potentially localized at the plasma membrane (Supplementary Fig. 5c, e). Because our tracking predominantly resolved PIEZO2 clusters rather than individual channels, these data indicate that STOML3 restricts the mobility of PIEZO2 clusters, consistent with a model where STOML3 either enriches local membrane cholesterol or constrains PIEZO channels within ring structures, or both.

### Ring formation and cholesterol contribute to efficient STOML3-channel interaction

To determine whether ring formation contributes to channel association, we used the P44S mutant that prevents large STOML3 ring assembly. Co-immunoprecipitation revealed that the P44S mutation differentially affected STOML3 interactions with mechanosensitive channels: interaction with PIEZO2 was reduced to approximately 29% of wild-type levels (a ∼70% reduction), whereas interaction with ELKIN1 was reduced to approximately 47% of wild-type levels (a ∼50% reduction) (Fig. 5a, b). An unexpected observation was that the T263N glycosylation mutant exhibited increased ring cluster intensity and a trend toward enhanced ELKIN1 interaction, although the latter did not reach statistical significance (Fig. 5a, b). Since this mutation does not alter membrane topology, N-glycosylation of the extracellular C-terminus may serve a regulatory rather than structural role—a possibility that warrants future investigation.

**Figure 5.**
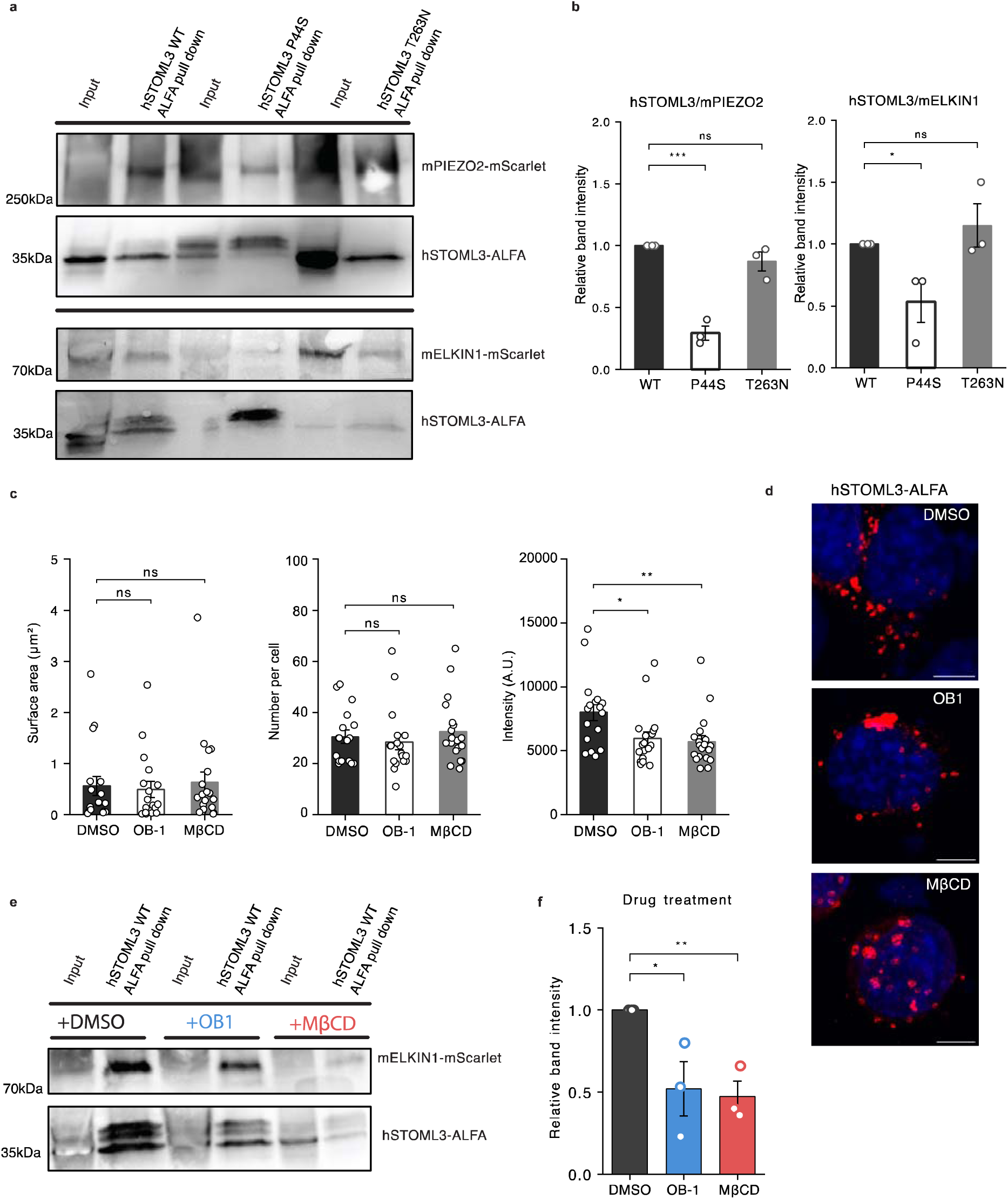
Ring formation and cholesterol contribute to efficient STOML3-channel interaction. **(a)** Co-immunoprecipitation of hSTOML3 variants with mPIEZO2-mScarlet and mELKIN1-mScarlet. ALFA-tag pull-down from cells expressing WT, P44S, or T263N hSTOML3-ALFA. Western blots show input and pull-down fractions for mPIEZO2-mScarlet (300 kDa), mELKIN1-mScarlet (70 kDa), and hSTOML3-ALFA (35 kDa). **(b)** Quantification of STOML3-PIEZO2 (left) and STOML3-mELKIN1 (right) interaction. P44S mutation reduces PIEZO2 interaction by approximately 70% (***P < 0.001) and ELKIN1 interaction by approximately 50% (*P < 0.05). T263N glycosylation mutant shows interaction levels comparable to WT (n.s., not significant). **(c)** Quantification of STOML3 cluster parameters following drug treatment. Cells expressing hSTOML3-ALFA were treated with DMSO (vehicle control), 10 μM OB-1 (STOML3 oligomerization inhibitor), or 5 mM MβCD (cholesterol depletion) and analyzed by Airyscan microscopy. Surface area and cluster numbers per cell are unchanged by either treatment (n.s.), whereas fluorescent intensity is reduced by both OB-1 (*P < 0.05) and MβCD (**P < 0.01), consistent with reduced STOML3 density within clusters despite maintained ring architecture. (n = 10 cells per condition from 3 independent experiments) **(d)** Representative Airyscan images of hSTOML3-ALFA (red) following treatment with DMSO, OB-1, or MβCD. Ring-like cluster morphology is preserved under all conditions. Scale bars, 5 μm. **(e)** Co-immunoprecipitation of hSTOML3-ALFA with mELKIN1-mScarlet following treatment with DMSO (vehicle), OB-1 (STOML3 inhibitor), or MβCD (cholesterol depletion). **(f)** Quantification showing both OB-1 (*P < 0.05) and MβCD (**P < 0.01) reduce STOML3-ELKIN1 interaction by approximately 50% compared to DMSO control. Data are mean ± SEM from n = 3 independent experiments. Statistical comparison by one-way ANOVA with Dunnett’s post-hoc test.

PIEZO2 and ELKIN1 show differential dependence on ring formation: ∼70% of PIEZO2 interaction requires intact oligomeric architecture, whereas ELKIN1 retains ∼50% of interaction with ring-deficient P44S-STOML3. This may indicate that ELKIN1 engages STOML3 through direct protein-protein contacts that persist when higher-order assembly is disrupted. Alternatively, the difference may reflect the much larger size of the PIEZO2 trimer compared with ELKIN1, which could make PIEZO2 more dependent on assembled rings.

To further dissect the ring-dependent component of channel interaction, we tested whether pharmacological or lipid perturbations phenocopy the P44S mutation. Treatment with OB-1, a small molecule inhibitor of STOML3 oligomerization^25^, reduced the STOML3-ELKIN1 interaction by approximately 50% compared to vehicle control (Fig. 5e, f). Cholesterol depletion using methyl-β-cyclodextrin (MβCD) produced a comparable ∼50% reduction in the STOML3-ELKIN1 interaction (Fig. 5e, f). Strikingly, the magnitude of reduction observed with OB-1 and MβCD treatment closely matches the ring-dependent fraction of ELKIN1 interaction defined by the P44S mutant (∼50%), consistent with these perturbations acting, at least in part, through disruption of ring assembly.

Neither OB-1 nor MβCD treatment altered STOML3 cluster surface area or cluster number (Fig. 5c, d), but both reduced cluster fluorescence intensity (Fig. 5c). This differs from the P44S mutation, which abolishes large rings and produces numerous small dispersed puncta (Fig. 2c, f, g). Ring-like clusters were therefore still present after both treatments, and neither compound disrupted the architecture of the rings that remained. The reduced intensity is consistent with a lower STOML3 content per cluster, which could reflect partial destabilisation of assembled rings, reduced incorporation of newly synthesised protein, or both. These experiments were performed in transiently transfected cells and report a single time point, and cannot distinguish between these possibilities.

## Discussion

Our data show that STOML3 adopts a predominantly Type II transmembrane topology, with a cytoplasmic N-terminus and an extracellular, N-glycosylated C-terminus. In contrast, for stomatin the hairpin-loop form appears to predominate and it is unclear what proportion is present as a transmembrane form^39,41,43,44^. Our results are consistent with work in oocytes, where it was found that approximately half of the STOML3 pool was N-glycosylated compared with a small minority for stomatin^44^. Both proteins carry the conserved membrane-proximal proline, so sequences outside this region must determine which configuration is favoured. Mouse and human STOML3 behaved identically, and the P40/P44 and T259/T263 positions are conserved between them. One consequence of the Type II arrangement is that the STOML3 C-terminus is exposed on the outer face of the membrane, where it is accessible to post-translational modification, an option not available to the cytoplasmic C-terminus of hairpin-configured stomatin. Whether glycosylation regulates STOML3 function remains open: the T263N mutant showed elevated cluster intensity but no significant change in ELKIN1 binding.

Different imaging modalities including Airyscan, STED and MINFLUX nanoscopy demonstrated that STOML3 assembled into ring-like structures at the plasma membrane considerably larger than any SPFH assembly described so far. STED gave a diameter distribution from 0.1 to 1.6 μm with a peak at 0.46 μm, and MINFLUX resolved rings of similar size (153–1,463 nm in diameter) containing 15 to almost 400 detected STOML3 molecules. Even the smallest of these rings was approximately an order of magnitude wider than the 16-mer stomatin dome^39,41,43^. The same structures appeared in N2a and HEK293T cells and in cultured DRG neurons, and were seen when the ALFA tag was labelled in living cells before fixation, arguing against a fixation or permeabilization artefact. The clusters were heterogeneous in size and in the number of detected STOML3 molecules, distinguishing them from the defined stoichiometry of purified SPFH complexes. The variable sizes that we observed have implications for what each assembly can enclose. The stomatin dome has an internal diameter of approximately 12 nm and a narrow intracellular opening, dimensions that accommodate small transmembrane proteins such as aquaporin-1 and urea transporter-B^41^ but would exclude a PIEZO trimer, whose curved blades form a dome of approximately 28 nm^53,54^.

STOML3 rings are one to two orders of magnitude larger and could in principle enclose several channel complexes together with the surrounding lipid. MINFLUX also revealed an axial extent of several hundred nanometres (median 328 nm), exceeding that of a single layer of membrane-bound STOML3. This could reflect STOML3-associated membrane deformation or a contribution from intracellular STOML3 labelled after permeabilization; surface-restricted labelling combined with membrane markers will be required to distinguish these possibilities. Whether STOML3 rings adopt a closed, dome-like architecture or a different organisation cannot be determined at the resolution used here. Their heterogeneity and size both make structural determination by single-particle cryo-EM very challenging, and cryo-electron tomography of native membranes may be better suited to this question.

We also showed that ring assembly depended on a conserved proline. The P44S mutation left STOML3 at the plasma membrane but redistributed it into approximately 224 small puncta per cell, in place of approximately 31 large clusters for the wild type protein. The equivalent residue in stomatin is required for hairpin topology and cholesterol binding^44,45^, and P40S mouse STOML3 fails to localize to lipid rafts and cannot rescue mechanosensitivity in STOML3-deficient neurons^31^. Our data now show that the conserved proline is necessary for STOML3 large ring assembly STOML3 co-fractionated with PIEZO2 and ELKIN1 in low-density membrane fractions, co-purified with PIEZO1, PIEZO2 and ELKIN1, and co-localised with all three channels by Airyscan microscopy, with Manders’ coefficients close to 0.9 against chance levels of 0.2 to 0.4. Our data are consistent with the idea that STOML3 rings might enclose a diverse group of mechanosensitive ion channels. However, the localization precision of our methods does not allow us to definitively conclude that STOML3 rings enclose the channels, this would still require atomic level measurements. In contrast to STOML3, stomatin showed neither co-purification nor co-localisation under the same conditions, so channel binding is not a general property of SPFH-domain proteins. Mass spectrometry identified endogenous ELKIN1 among the most enriched proteins, supporting the interaction at native levels. Live TIRF imaging showed that PIEZO2 clusters move together with STOML3 clusters, and that the diffusion coefficients of sub-diffusive and confined PIEZO2 cluster populations are higher in cells lacking STOML3, an effect comparable to that of cholesterol depletion on PIEZO1 mobility in other systems^55,56^. Together these observations indicate that STOML3 restricts the lateral mobility of PIEZO2 clusters, although our experiments do not establish whether this reflects confinement within rings, a change in the local lipid environment, or both. Perturbing ring assembly consistently reduced channel co-immunoprecipitation but never abolished it. P44S-STOML3 retained approximately 29% of PIEZO2 and 47% of ELKIN1 binding relative to wild-type, and OB-1 and MβCD produced comparable reductions in ELKIN1 binding. One explanation is that STOML3 engages these channels in two ways, through an association that does not require assembled rings and a further component that depends on them. A second is that the residual association is not a direct STOML3–channel contact but is bridged by other proteins that remain in the same membrane domain when ring assembly fails; co-immunoprecipitation reports proximity within a complex rather than direct binding. PIEZO2 was consistently more sensitive to ring disruption than ELKIN1, which may reflect differences in channel architecture or a greater reliance of ELKIN1 on such bridging interactions.

Our interactome offers candidates for such partners, and raises the further possibility of mixed assemblies. Endogenous stomatin was the eighth most enriched protein in the STOML3 interactome (log_₂_ fold-change = 5.1), and the mitochondrial family member STOML2 was also recovered (log_₂_ fold-change = 2.2). Hetero-oligomerisation recurs across the SPFH superfamily: prohibitins form obligate PHB1–PHB2 rings, and HflK and HflC assemble into a heteromeric cage^57,58^. Since stomatin does not itself bind these channels, its enrichment more likely reflects a shared membrane domain, or co-assembly with STOML3, than a shared channel complex. Mixed subunit composition would also account for the heterogeneity in ring dimensions described above, and whether composition tunes ring size or channel selectivity is now directly testable.

How ring assembly alters channel behaviour remains to be established. MβCD treatment strongly attenuates STOML3-dependent potentiation of mechanosensitivity^31^, and the membrane-stiffening mechanism proposed previously^31^ is consistent with our data. Thus, large STOML3 rings may act by altering the mechanical properties of the surrounded lipid bilayer rather than facilitating gating of enclosed ion channels. This would also explain why cryo-EM structures of stomatin do not resolve bound cholesterol despite clear biochemical evidence for cholesterol dependence^38^, since cholesterol may act on SPFH assemblies through the lipid environment rather than through a defined binding pocket.

Among the SPFH proteins, STOML3 occupies an unusual position: it is largely restricted to sensory neurons and is so far the only family member for which pharmacological inhibition of oligomerisation has produced a therapeutic effect in vivo^25^. The results presented here will allow a better understanding of how STOML3-drug interactions may work. STOML3 does not function as an isolated oligomer but as a large, cholesterol-dependent ring that concentrates PIEZO1, PIEZO2 and ELKIN1 at the plasma membrane. Establishing the subunit stoichiometry of these rings and the interfaces that hold them together will require atomic-resolution structures of STOML3 alone and in complex with its channel partners, and is the next step towards targeting the assembly rationally.

## Methods

### Animals

All experiments involving animals were performed in accordance with German federal and European Union legislation on the protection of animals used for scientific purposes. Mice were housed in groups under a 12 h light/dark cycle with food and water available ad libitum. Dorsal root ganglia were obtained from adult C57BL/6J mice. All animal protocols were approved by the German federal authorities (State of Berlin). Before the dissection of dorsal root ganglia, all animals were euthanized by cervical dislocation.

### Cell culture and transfection

HEK293T cells (ATCC, CRL-3216) were maintained in Dulbecco’s Modified Eagle Medium (DMEM; Gibco, 41966029) supplemented with 10% fetal bovine serum (FBS; Gibco) and 1% penicillin-streptomycin (Gibco, 15140122) at 37°C in a humidified atmosphere containing 5% CO₂. N2a (Neuro-2a) cells, N2a-STOML3KO cells, and N2a-PIEZO1KO cells were maintained in DMEM supplemented with 45% Opti-MEM (Gibco), 10% FBS, and 1% penicillin-streptomycin under the same conditions. N2a-STOML3KO cells were generated by CRISPR/Cas9 gene editing using guide RNA sequences targeting exon of STOML3 as previously described^52^. For transfection, cells were seeded at 2×10 cells/well in 12-well plates and transfected at 70% confluency using FuGENE HD (4 μL per 100 μL Opti-MEM with 1 μg cDNA) according to the manufacturer’s instructions. Cells were analyzed 48 hours post-transfection.

### Plasmid construction

Mouse STOML3 (mSTOML3; UniProt: Q6PE84) and human STOML3 (hSTOML3; UniProt: Q8TAV4) coding sequences were cloned into pRK8 using Gibson assembly.

### ALFA-tagged constructs

The ALFA-tag sequence (SRLEEELRRRLTE) was fused to the C-terminus of STOML3 to generate mSTOML3-ALFA and hSTOML3-ALFA constructs, enabling detection of the extracellular C-terminus with anti-ALFA nanobodies.

### Tripartite split-GFP topology reporters

GFP10 and GFP11 fragments were fused to either the N- or C-terminus of mSTOML3. Co-expression indicators (RFP for GFP10-tagged, BFP for GFP11-tagged constructs) were included via IRES elements for expression verification^6^.

### Point mutants

Glycosylation site mutants (mSTOML3-T259N, hSTOML3-T263N) and cholesterol-binding site mutants (mSTOML3-P40S, hSTOML3-P44S) were generated by site-directed mutagenesis using the Q5 Site-Directed Mutagenesis Kit (NEB, E0554S). Human stomatin (hSTOM-ALFA) was cloned similarly as a negative control. The N-glycosylation consensus-site mutants used for Western blot analysis (S18N, S224N and T259N; Fig. 1b) were generated by PCR-based site-directed mutagenesis in a 3xFLAG-tagged mSTOML3 background.

### Channel constructs

Mouse PIEZO1-mScarlet and PIEZO2-mScarlet were gifts from Prof. Dr. Stefan Lechner (UKE). Mouse ELKIN1-mScarlet was generated by Gibson assembly. All constructs were verified by Sanger sequencing (LGC Genomics). To generate PIEZO2-HaloTag, the coding sequence of HaloTag was inserted at mouse PIEZO2-Cter with a GSGS linker using Gibson assembly (NEBuilder HiFi, New England Biolabs). DNA fragments were amplified by PCR (primers from Sigma) using KAPA HiFi polymerase (Roche), digested with DpnI (New England Biolabs) and column purified with standard kit (PureLink, Invitrogen). After homologous recombination, mixture was transformed into electrocompetent Stbl4 bacteria (Invitrogen) and grown at 30°C for 48 hours. Selected clones were entirely sequenced (Eurofins) to ensure proper insertion and that no mutations were present.

### AAV transduction of DRG neurons

DRG from adult C57BL/6J mice, dissociated with collagenase, trypsin for 1 hour and 15 mins, plated on PLL and iMatrix coated cover slip in F12/horse serum medium. Transduced with AAV-PHP.S-hSyn-mSTOML3-ALFA or AAV-PHP.S-tdTomato at 5*10^10^(VG/ml) for 24 hours after plating; imaged 6 days later. Surface staining and Airyscan as above.

### Sucrose gradient fractionation

Lipid raft fractionation was performed using discontinuous sucrose density gradient centrifugation. Cells were lysed in 1 mL ice-cold HBS buffer (25 mM HEPES pH 7.5, 150 mM NaCl) containing 1% Triton X-100 and protease inhibitors. Lysates were incubated on ice for 5 minutes and homogenized. Cell lysates were adjusted to 40% sucrose by mixing with an equal volume of 80% sucrose in HBS buffer. The lysate (2 mL) was placed at the bottom of an ultracentrifuge tube and overlaid with 5 mL of 35% sucrose and 3.5 mL of 5% sucrose in HBS buffer. Gradients were centrifuged at 200,000 × g for 18 hours at 4°C in a 70.1Ti rotor. Thirteen fractions of 1 mL each were collected from the top of the gradient. Fractions were analyzed by Western blot. Flotillin-2 (Sigma, F1805) served as a lipid raft marker. Low-density fractions (fractions 2–5) contain lipid rafts, while high-density fractions contain non-raft membranes.

### Tripartite split-GFP topology assay

The tripartite split-GFP assay was used to determine STOML3 membrane topology^59,60^. The system consists of three GFP fragments: GFP1-9 (detector), GFP10, and GFP11. GFP fluorescence is reconstituted only when all three fragments associate in the same cellular compartment.

**Construct combinations tested:**

• N-N: GFP10-mSTOML3 + GFP11-mSTOML3 (both N-termini tagged)

• N-C: GFP10-mSTOML3 + mSTOML3-GFP11 (N- and C-termini tagged)

• C-C: mSTOML3-GFP10 + mSTOML3-GFP11 (both C-termini tagged)

• Vector control: GFP10-mSTOML3 or mSTOML3-GFP10 + GFP11 vectors

GFP1-9 detector was co-transfected in HEK293T cells. Co-expression was verified by BFP fluorescence. Cells were imaged 48 hours post-transfection using a BioTek Cytation plate reader equipped with DAPI/BFP, GFP, and RFP LED/filter cubes. GFP-positive cells were quantified from 4× fields of view per condition across 3 independent experiments. Percentage of GFP-positive cells among double-positive (RFP⁺BFP⁺) cells was calculated.

### Western blotting and glycosylation analysis

#### Sample preparation

Cells were lysed in RIPA buffer (50 mM Tris-HCl pH 7.4, 150 mM NaCl, 1% NP-40, 0.5% sodium deoxycholate, 0.1% SDS) supplemented with protease inhibitor cocktail (Roche, A154.2). Protein concentrations were determined by Bradford assay (Bio-Rad, 5000006).

### Deglycosylation

Lysates were divided into two equal aliquots and treated with PNGase F (Roche) essentially as described for stomatin^44^. Samples were dissolved in denaturing buffer (0.1% SDS, 50 mM 2-mercaptoethanol) and incubated at 56 °C for 10 min. Sodium phosphate (pH 8.0) was then added to a final concentration of 80 mM and samples were incubated for 3 h at 37 °C with PNGase F in the presence of 1% (v/v) Triton X-100. Control aliquots were treated identically but without PNGase F. Both aliquots were supplemented with gel loading buffer and analysed in parallel by SDS-PAGE and Western blotting as described below.

### Gel electrophoresis

Equal amounts of protein (∼10 μg) were separated on 10% SDS-PAGE gels and transferred to nitrocellulose membranes (Fisher Scientific, 15299794) at 220 V for 90 minutes.

### Immunoblotting

Membranes were blocked in 3% non-fat milk in TBST for 20 minutes at room temperature and incubated overnight at 4°C with primary antibodies: anti-ALFA nanobody-HRP (NanoTag Biotechnologies, N1505, 1:500), anti-Flotillin-2 (Sigma, F1805, 1:1000), anti-RFP (Rockland, 36970, 1:500), or anti-FLAG M2-HRP (Sigma-Aldrich, 1:8000). After washing, membranes were incubated with HRP-conjugated secondary antibodies (1:1000) for 1 hour at room temperature. Bands were detected using ECL substrate (Thermo Fisher, 34580) and imaged on a Bio-Rad imaging system.

### Immunocytochemistry

Cells were seeded on poly-L-lysine-coated glass coverslips in 12-well plates and transfected as described above.

### Surface staining (non-permeabilized)

Cells were washed with ice-cold PBS and fixed with 4% paraformaldehyde in PBS for 10 minutes at room temperature. Cells were NOT permeabilized to ensure detection of only surface-exposed ALFA-tag, providing independent validation of extracellular C-terminus topology. Cells were blocked in 10% goat serum in PBS for 15 minutes, then incubated with anti-ALFA nanobody conjugated to Star635P or IRDye 680RD (NanoTag Biotechnologies, N1502-Ab635P-L or N1502-Li680-L, 1:500) for 1 hour at room temperature. After washing 3 times with PBS containing DAPI (1:1000), coverslips were mounted and cured overnight before imaging.

For colocalization studies with mScarlet-tagged channels, the same non-permeabilized staining protocol was used; mScarlet fluorescence was directly imaged without additional antibody staining.

### Live-cell surface labelling

live N2a cells or DRG neurons incubated with anti-ALFA nanobody-Star635P (1:500) in DMEM medium for 1 hour at 37°C, washed 3 times with PBS, then fixed in 4% PFA for 10 min without permeabilization.

### Airyscan microscopy

Airyscan imaging was performed on a Zeiss LSM 980 confocal microscope (Axio Observer.Z1/7) equipped with an Airyscan 2 detector module.

**Imaging parameters:**

• Objective: C Plan-Apochromat 63×/1.40 Oil

• Laser lines: 639 nm for Star635P (single expression); 561 nm for mScarlet and 680 nm for IRDye 680RD (co-expression)

• Airyscan mode: SR (super-resolution)

• Pixel size: 35 nm × 35 nm

• Frame averaging: 4

• Image size: 2930 × 2930 pixels (103.39 × 103.39 μm)

• Bit depth: 16-bit

### Image processing

Raw Airyscan images were processed using Zeiss ZEN software with the Airyscan processing module (3D processing, automatic strength). Theoretical resolution: ∼120 nm laterally and ∼350 nm axially.

### STED microscopy

STED (Stimulated Emission Depletion) microscopy was performed on a Leica TCS SP8 STED 3X system.

### Sample preparation

Cells were stained with anti-ALFA nanobody conjugated to Star635P for STOML3 detection.

**Imaging parameters:**

• Objective: HC PL APO 100×/1.40 Oil STED WHITE

• Excitation: 633 nm

• Depletion laser: 775 nm pulsed

• Emission detection: 650–750 nm

• Pixel size: 20 nm × 20 nm

• Scan speed: 100 Hz

• Frame averaging: 3

• Image format: 2048 × 2048 pixels

• Zoom: 4.5×

### Image processing

Images were deconvolved using Leica LAS X software. Lateral resolution achieved: ∼40–60 nm. Depletion was applied in the lateral plane only, so axial resolution remained close to that of confocal microscopy (∼500–600 nm). Ring diameter measurements were performed on 255 rings from 9 cells across 3 independent experiments.

### Tau-STED microscopy

Representative Tau-STED images (Fig. 2i, j) were acquired on the same Leica TCS SP8 STED 3X system using time-correlated single photon counting (TCSPC)-based fluorescence lifetime gating (TauSTED mode in Leica LAS X software). Imaging parameters were identical to standard STED acquisition (see above), with the addition of lifetime-based separation of depleted and non-depleted fluorophore populations to improve effective resolution and signal-to-noise ratio at reduced depletion laser power. Confocal reference images were acquired simultaneously from the same scan. DAPI was excited at 405 nm.

### MINFLUX nanoscopy

N2a cells were plated on PLL-coated 18mm coverslips as described in previous section and transfected with N-terminally GFP-tagged hSTOML3. Two to three days after transfection, cells were washed once with PBS and fixed for 10min at room temperature with a mixture of 1% PFA and 0.05% glutaraldehyde in PBS. Fixative was removed and quenched with 5mM NaBH_4_ in PBS (one quick wash and another one for 5min at room temperature) and then with 50mM glycine and 50mM NH_4_Cl in PBS (one quick wash and another one for 5min at room temperature). Samples were further washed thrice with PBS for 5min at room temperature, and were blocked with Antibody Incubation buffer (Massive Photonics; contains Tween) for 30min at room temperature. The cells were incubated overnight at 4°C with Massive-Tag-Q Anti-GFP nanobody conjugated with a DNA docking strand (Massive Photonics) diluted at 1:200 (Anti-GFP) in Antibody Incubation buffer. Cells were then washed thrice with 1X Washing buffer (Massive Photonics) and then post-fixed for 5min. Fixatives were quenched and washed as described above. Samples were then incubated for 10 min with 100-200 µl of gold nanoparticles for future stabilization (gold colloid 250nm, BBI Solutions). Unbound nanoparticles were rinsed extensively with PBS and remaining nanoparticles were further stabilized with PLL for at least 1hr at room temperature. Cells were then washed thrice with PBS before mounting. Labelled samples were used within 2 days. DNA-PAINT imagers (Massive Photonics) conjugated to Atto 655 were freshly diluted in Imaging buffer (Massive Photonics) for a final concentration of 1nM. A drop of imager dilution was added into a cavity slide and coverslips were mounted and sealed with picodent twinsil (picodent).

MINFLUX imaging was performed on an Abberior MINFLUX commercial microscope using an inverted IX83 microscope with a 100X UPlanXApo objective (Olympus) and with Imspector Software (Abberior Instruments). Routine alignment and calibration of the excitation beam pattern and position of the pinhole was performed using fluorescent nanoparticles (abberior Nanoparticles, Gold 150 nm, 2C Fluor 120 nm, Abberior Instruments). Cells were identified with a 488nm (hSTOML3-GFP) confocal scan and the transient binding of imagers with Atto655 was quickly verified with quick 640 nm confocal scan. At least 4 gold fiducials were present in the field of view and used by the active-feedback stabilization system of the microscope (IR 975nm laser, Cobolt, and CCD camera, The Imaging Source), yielding a precision below 1nm in all three axes and being stabled for hours. Several ROIs of 1x1 to 4x4 µm having STOML3 rings were selected at the cell-coverslip interface. Laser power in the first iteration and pinhole was set at 16% power and 0.83 A.U pinhole. Final laser power in the last iteration is scaled up by a factor of six. ROIs were imaged for at least 2 hours and up to overnight using the standard 3D MINFLUX sequence. Detection for Atto 655 signals was performed with two avalanche photodiodes channels (650-685nm and 685-720nm) that were added. Specificity of the nanobodies and imagers used in this study and of the MINFLUX signal was evaluated before^52^.

Final valid localizations were exported from Imspector software as matlab files and custom Matlab scripts were then used for all subsequent operation. Data filtering involved first an efo filter (effective frequency at offset, kHz, retrieved for each individual valid locations) based on its overall distribution for each individual measurements: a threshold was then selected to filter-out potential multiple emitters. Then, localizations from the same emission trace, i.e. with the same trace identification number (TID), having a standard deviation of more than 10nm in the x, y, z axes and having less than 3 localizations were excluded. Localizations for each trace were aggregated (group of 3), as described elsewhere^61^. Remaining traces were further corrected for the refractive index mismatch between the coverslip and the sample, applying a scaling factor of 0.7 for all traces in the z dimension^62^. The resulting localization precision was ∼3 nm in x, y and z (Fig. 3d).

A density-based clustering (DBSCAN) step was used on the raw STOML3 filtered localizations to identify clusters of molecules, starting with a minimal number of localizations (minPoints) of 100 and a search radius (epsilon) of 100nm and adjusted depending on the signal density, followed by a manual inspection and refinement. Then, the center of mass of each individual filtered trace (i.e trace with the same TID) within the clusters was calculated. Due to the nature of DNA-PAINT labelling and the possible proximity of individual GFP molecules at STOML3 N-ter, neighbouring signals coming from potentially the same STOML3 protein were estimated and averaged using another DBSCAN on the trace average calculated before, with a minPoints of 2 and epsilon of 8nm (based on recording precision and labelling error), ultimately giving a position of an individual STOML3 protein. Identified clusters were then processed individually to measure their dimensions. For each cluster, a centre point was determined from the averaged traces within the lowest third of the cluster height, and the radius was measured as the distance from this centre to the outermost trace; ring diameter is reported as twice this radius. Axial extent was calculated as the difference between the lowest and highest traces of the cluster. Nearest-neighbour distances were computed between the resulting STOML3 positions in MATLAB.

### Live-cell TIRF microscopy

N2a-PIEZO1KO and N2a-STOML3KO cells were prepared as described above. Cells were seeded on PLL-coated 35mm glass-bottom dishes (MatTek High Precision #1.5 Coverslip) and transfected the next day with PIEZO2-HaloTag plasmid. Medium was changed the next day and replaced by phenol red-free one. Two to three days post-transfection, cells were labelled with JFX-650 Halo ligand (Promega) the day before imaging (1nM, overnight incubation), and rinsed for at least 20 minutes with medium before imaging. TIRF imaging was performed on a Nikon Eclipse TiE microscope, with a Roper iLAS2 TIRF module. A Nikon CFI Plan Apo Lambda 100x (NA 1.45) objective was used, together with a Photometrics Prime 95B back-illuminated sCMOS camera, having a resolution of 1200 x 1200 and with a pixel size of 11 µm. A 1.5x magnification lens was added, giving a final pixel size of 0.073µm. Cells were illuminated with a solid-state laser of 640nm (20% power). Acquisition was done with VisiView software (version 5.0.0, Visitron Systems). An incubation chamber (okolab) was used to adjust temperature (37°C), CO2-concentration (5%) and humidity. Live cells were imaged for 30s with a frame rate of 10Hz (approximately 100ms exposure time per frame).

### Single-particle tracking analysis

Clusters and tracks analysis from live-imaging were processed as described before^63,64^. PIEZO track analysis was performed with TrackMate v7.13.2^65,66^ and the following parameters: DoG detector, blob diameter of 0.35µm, spots quality filter value of 0.30-2, simple LAP tracker with a linking distance of 0.3µm, a gap closing distance of 0.3µm and a maximal gap closing frame number of 2. Only tracks that have a duration of at least 40 frames were considered for further analysis. Tracks classification was done in ImageJ with the TraJClassifier plugin^67^ and the following parameters: minimal track length 40, window size 30, minimal segment length 30. MSD calculation and subsequent analysis were then performed in Matlab (Mathworks) using custom scripts (available at: https://github.com/clementverkest/PIEZO_Track_Analysis).

### Affinity purification and co-immunoprecipitation

For co-immunoprecipitation, HEK293T cells were co-transfected with STOML3-ALFA constructs and mScarlet-tagged channel constructs (PIEZO1, PIEZO2, or ELKIN1).

### Cell lysis

48 hours post-transfection, cells were washed with ice-cold PBS and lysed in IP buffer (50 mM Tris-HCl pH 7.4, 150 mM NaCl, 1% NP-40, 0.5% sodium deoxycholate, 0.1% SDS) supplemented with protease inhibitors. Lysates were incubated on ice for 5 minutes, sonicated (10 seconds on, 40 seconds off, 4 cycles), and clarified by centrifugation at maximum speed for 25 minutes at 4°C.

### ALFA-tag affinity purification

Clarified lysates were incubated with ALFA-SelectorPE beads (NanoTag Biotechnologies, N1510) at 20 μL beads per 500 μL lysate for 1 hour at 4°C with rotation. Beads were washed 3 times with IP buffer. Bound proteins were eluted by competitive elution with ALFA peptide (100 μM) at 4°C for 30 minutes.

Samples were analyzed by Western blot using anti-ALFA and anti-RFP antibodies. Band intensities were quantified using iBright analysis software and normalized to pulled-down STOML3 levels.

### Mass spectrometry

#### Sample preparation and digestion

STOML3-ALFA was immunoprecipitated from HEK293T cells as described above. Cells expressing the ALFA tag alone served as a control for immunoprecipitation. Bound proteins were eluted by competitive elution with 100 μM ALFA peptide for 30 min at 4 °C. Eluates from three biological replicates each of STOML3-ALFA and control immunoprecipitations were digested using a modified SP3 protocol^68^. Briefly, proteins were reduced with 5 mM TCEP and alkylated with 40 mM chloroacetamide (CAA) for 1 h at room temperature in the dark. A 1:1 (v/v) mixture of hydrophilic and hydrophobic Sera-Mag beads was added, followed by acetonitrile (ACN) to a final concentration of 50%. After removal of the supernatant, the beads were washed twice with ethanol and once with ACN. Proteins were digested on-bead in 50 mM triethylammonium bicarbonate (TEAB, pH 8.5) with Lys-C and trypsin, each at an enzyme-to-protein ratio of 1:50 (w/w), for 16 h at 37 °C. The beads were subsequently washed twice with ACN to a final concentration of >95%. Peptides were eluted with 5% DMSO and dried for subsequent analysis.

### LC-MS/MS analysis

Peptides were resuspended in 1% ACN with 0.05% trifluoroacetic acid (TFA). For each run, 500 ng of peptides were injected into a Vanquish Neo UHPLC system (Thermo Fisher Scientific) connected to a PepMap C18 trap column (75 µm × 50 mm, 3 µm particle size, 100 Å pore size; Thermo Fisher Scientific), followed by an in-house packed C18 analytical column (Poroshell 120 EC-C18, 2.7 µm; Agilent Technologies). Peptides were separated at a flow rate of 250 nL/min using a 120-min gradient of increasing ACN concentration and analysed on an Orbitrap Exploris 480 mass spectrometer (Thermo Fisher Scientific; Instrument Control Software version 4.2) in data-dependent acquisition (DDA) mode. MS1 scans were acquired in the Orbitrap at a resolution of 120,000 over m/z 375–1,200, with 50% RF lens, 300% AGC target and automatic maximum injection time. MS2 scans were acquired in the Orbitrap with automatic scan range, 100% AGC target, automatic maximum injection time, an isolation window of 1.6 m/z, a normalized collision energy (NCE) of 30% and a resolution of 15,000. Only precursors with charge states +2 to +4 were fragmented, and fragmented precursors were dynamically excluded for 20 s. Data were acquired in 2-s cycles.

### Protein identification and label-free quantification

DDA data were analysed using FragPipe v23.0 with MSFragger v4.2^69^, Philosopher v5.1.1^70^ and IonQuant v1.11.9^71^. Data were searched against the human proteome (one protein per gene, retrieved on 5 May 2025) supplemented with common contaminants and reverse decoy sequences. The LFQ-MBR workflow in FragPipe was used with default settings unless stated otherwise: precursor tolerance ±10 ppm, fragment tolerance ±20 ppm, mass recalibration and parameter optimization activated, trypsin cleavage C-terminal to K/R with up to two missed cleavages, peptide length 7–50 residues, peptide mass 500–5,000 Da, oxidation of methionine and acetylation of protein N-termini as variable modifications, and carbamidomethylation of cysteine as a static modification.

### Statistical analysis

Enrichment analysis was performed in R using the limma package^72^. Contaminants and proteins with fewer than two of three values in both conditions were removed. Data were median-normalized, and missing values were imputed by random sampling from a downshifted normal distribution (downshift 1.8, width 0.3), following the standard Perseus imputation approach. Proteins with P < 0.02 and |log fold-change| > 1 were considered significantly enriched.

### Gene Ontology analysis

GO enrichment of the 1,393 significantly enriched proteins was performed with g:Profiler (g:GOSt, Homo sapiens; accessed 31 March 2026)^73^, using all annotated human genes as the statistical background and g:SCS correction for multiple testing (adjusted P < 0.05). g:GOSt calculates enrichment P values with a one-sided hypergeometric test, equivalent to a one-sided Fisher’s exact test. GO terms annotated to more than 2,000 genes were excluded and redundant parent–child terms were removed; ten representative terms per category (biological process, cellular component and molecular function) are shown in Supplementary Fig. 4c.

### Drug treatments

#### OB-1 treatment

OB-1 (small-molecule inhibitor of STOML3 oligomerization) was dissolved in DMSO. Cells were treated with 10 μM OB-1 or equivalent volume of DMSO (vehicle control) for 24 hours at 37°C prior to analysis.

#### Cholesterol depletion

Methyl-β-cyclodextrin (MβCD; Sigma-Aldrich, C4555) was dissolved in serum-free medium. Cells were treated with 5 mM MβCD for 2–3 hours at 37°C^31^. For imaging experiments, cells were treated, then fixed and stained as described. For co-immunoprecipitation, cells were treated, then lysed and processed immediately.

### Cluster analysis

Image analysis was performed using FIJI/ImageJ^74^. Background subtraction was performed using the rolling ball algorithm (radius: 50 pixels). Images were thresholded using the Otsu method. Cluster analysis was performed using the Analyze Particles function with a minimum size threshold of 0.01 μm².

**Parameters measured per cell:**

• Surface area: Average of all cluster areas (μm²)

• Cluster number: Count of discrete clusters per cell

• Fluorescent intensity: Mean integrated density per cluster, background-corrected

Quantification was performed on maximum intensity projections of 50–80 Z-slices covering the plasma membrane. At least 10 cells were analyzed per condition from 3 independent experiments.

### Ring diameter measurement (STED)

Ring diameters were measured as the full-width at half-maximum (FWHM) of line profiles drawn across ring centers, measured from the longest axis of each ring structure.

### Colocalization analysis

Colocalization analysis was performed using the ZEN 3.9 (Zeiss) Coloc module. Channel alignment was verified using TetraSpeck beads. Costes automatic thresholding was applied.

### Colocalization metrics

Manders’ M1 coefficient was calculated as the fraction of STOML3 signal overlapping with channel signal. Statistical validation was performed using the Costes randomization test with 3 iterations. Chance colocalization levels were determined by pixel randomization.

Analysis was performed on more than 10 cells per condition from 3 independent experiments.

### Sequence alignment

Protein sequences were retrieved from UniProt database: *C. elegans* MEC-2 (Q27433), mouse STOML3 (Q6PE84), and human STOML3 (Q8TAV4). Multiple sequence alignment was performed using Clustal Omega with default parameters. Alignments were visualized using custom Python scripts (matplotlib 3.7.1). Key conserved features annotated include the membrane-proximal proline (P40 mSTOML3, P44 hSTOML3) and N-glycosylation site (T259 mSTOML3, T263 hSTOML3).

### Statistical analysis

Statistical analyses were performed using GraphPad Prism version 10. Data are presented as mean ± SEM. Individual data points are shown in all graphs. Sample size n represents number of cells analyzed from N = 3 independent experiments.

Normality testing was assessed by Shapiro-Wilk test. Two-group comparisons used unpaired two-tailed Student’s t-test (normal distribution) or Mann-Whitney U-test (non-normal distribution). Multiple group comparisons used one-way ANOVA with appropriate post-hoc tests for normally distributed data, or Kruskal-Wallis with Dunn’s post-hoc test for non-normal data.

Significance thresholds: *P < 0.05, **P < 0.01, ***P < 0.001, ****P < 0.0001; n.s., not significant (P ≥ 0.05). Exact P-values are reported in figure legends or source data.

## Data availability

Source data for all figures are provided with this paper. Mass spectrometry data including identified proteins and statistical analyses are provided in Supplementary Table 1. The mass spectrometry proteomics data have been deposited to the ProteomeXchange Consortium via the PRIDE partner repository^75^ with the dataset identifier PXD087379. All other data supporting the findings of this study are available from the corresponding author upon reasonable request.

## Code availability

Custom Matlab scripts for PIEZO track analysis are available at: https://github.com/clementverkest/PIEZO_Track_Analysis

## Acknowledgements

We thank Dr. Alice Rossi, Franziska Bartelt, Kathleen Barda for technical assistance and Prof. Dr. Oliver Daumke for helpful discussions. We thank the Advanced Light Microscopy Facility at Max Delbruck Center Mitte and Leica confocal specialist Andreas Fulterer for assistance with STED imaging. We thank Dr. Antonio Failla and the team of the imaging core facility at UKE Hamburg (DFG Research Infrastructure Portal #RI_00489) for technical assistance with the MINFLUX and TIRF microscope. We thank Heike Stephanowitz (Mass Spectrometry Core Facility, FMP) for proteomics support. This work was supported by the European Research Council (https://erc.europa.eu/); ERC-2018-ADG-789128 and ERC-2023-ADG-101142488 to GRL and Postdoctoral Research Abroad Program sponsored by Ministry of Science and Technology (Taiwan) grant number 111-2917-I-564-011 to A.T.-L.H during 2022-2023. Funding for the MINFLUX microscope was awarded by the Hamburgische Investitions-und Förderbank (IFB, grant no. 51164232) under the Operational Programme Hamburg ERDF 2014– 2020, REACT-EU of the European Regional Development Fund (ERDF).

## Author contributions

A.T.-L.H. conceived and designed the study, performed experiments, analyzed data, and wrote the manuscript. J.H. performed the N-glycosylation site mutagenesis and the PNGase F deglycosylation experiments. M.R. and F.L. performed mass spectrometry and analysed proteomics data. C.V. and S.G.L. performed MINFLUX, single-particle tracking, live TIRF experiments, analysed the data, and provided channel constructs. G.R.L. co-conceived and supervised the project, secured funding, and edited the manuscript. All authors reviewed and approved the final manuscript.

## Competing interests

The authors declare no competing interests.

## Notes

### Competing Interest Statement

The authors have declared no competing interest.

## References

1. Katta, S., Krieg, M. & Goodman, M. B. Feeling force: physical and physiological principles enabling sensory mechanotransduction. Annu Rev Cell Dev Biol 31, 347–71 (2015).

2. O’Hagan, R., Chalfie, M. & Goodman, M. B. The MEC-4 DEG/ENaC channel of Caenorhabditis elegans touch receptor neurons transduces mechanical signals. Nat. Neurosci. 8, 43–50 (2005).

3. Yan, Z. et al. Drosophila NOMPC is a mechanotransduction channel subunit for gentle-touch sensation. Nature 493, 221–225 (2013).

4. Coste, B. et al. Piezo1 and Piezo2 Are Essential Components of Distinct Mechanically Activated Cation Channels. Science 330, 55–60 (2010).

5. Ranade, S. S. et al. Piezo2 is the major transducer of mechanical forces for touch sensation in mice. Nature 516, 121–U330 (2014).

6. Chakrabarti, S. et al. Touch sensation requires the mechanically gated ion channel ELKIN1. Science 383, 992–998 (2024).

7. Chalfie, M. Neurosensory mechanotransduction. Nat Rev Mol Cell Biol 10, 44–52 (2009).

8. Hu, J., Chiang, L. Y., Koch, M. & Lewin, G. R. Evidence for a protein tether involved in somatic touch. Embo J. 29, 855–67 (2010).

9. Brown, A. L., Liao, Z. & Goodman, M. B. MEC-2 and MEC-6 in the *Caenorhabditis elegans* Sensory Mechanotransduction Complex: Auxiliary Subunits that Enable Channel Activity. J. Gen. Physiol. 131, 605–616 (2008).

10. Das, A. et al. C. elegans touch receptor neurons direct mechanosensory complex organization via repurposing conserved basal lamina proteins. Curr. Biol. CB 34, 3133–3151.e10 (2024).

11. Chalfie, M. & Au, M. Genetic Control of Differentiation of the *Caenorhabditis elegans* Touch Receptor Neurons. Science 243, 1027–1033 (1989).

12. Ernstrom, G. G. & Chalfie, M. Genetics of Sensory Mechanotransduction. Annu. Rev. Genet. 36, 411–453 (2002).

13. Syntichaki, P. & Tavernarakis, N. Genetic Models of Mechanotransduction: The Nematode *Caenorhabditis elegans*. Physiol. Rev. 84, 1097–1153 (2004).

14. Huang, M., Gu, G., Ferguson, E. L. & Chalfie, M. A stomatin-like protein necessary for mechanosensation in C. elegans. Nature 378, 292–295 (1995).

15. Tavernarakis, N., Driscoll, M. & Kyrpides, N. C. The SPFH domain: implicated in regulating targeted protein turnover in stomatins and other membrane-associated proteins. Trends Biochem. Sci. 24, 425–427 (1999).

16. Ma, C. et al. Structural insights into the membrane microdomain organization by SPFH family proteins. Cell Res. 32, 176–189 (2022).

17. Daumke, O. & Lewin, G. R. SPFH protein cage — one ring to rule them all. Cell Res. 32, 117–118 (2022).

18. Lapatsina, L., Brand, J., Poole, K., Daumke, O. & Lewin, G. R. Stomatin-domain proteins. Eur. J. Cell Biol. 91, 240–245 (2012).

19. Fricke, B. et al. The ‘stomatin’ gene and protein in overhydrated hereditary stomatocytosis. Blood 102, 2268–2277 (2003).

20. Martinez-Salgado, C. et al. Stomatin and sensory neuron mechanotransduction. J. Neurophysiol. 98, 3802–3808 (2007).

21. Zhu, Y. et al. Stomatocytosis is absent in ‘stomatin’-deficient murine red blood cells. Blood 93, 2404–2410 (1999).

22. Wang, D., Mentzer, W. C., Cameron, T. & Johnson, R. M. Purification of band 7.2b, a 31-kDa integral phosphoprotein absent in hereditary stomatocytosis. J. Biol. Chem. 266, 17826– 17831 (1991).

23. Hiebl-Dirschmied, C. M., Adolf, G. R. & Prohaska, R. Isolation and partial characterization of the human erythrocyte band 7 integral membrane protein. Biochim. Biophys. Acta BBA - Biomembr. 1065, 195–202 (1991).

24. Wetzel, C. et al. A stomatin-domain protein essential for touch sensation in the mouse. Nature 445, 206–209 (2007).

25. Wetzel, C. et al. Small-molecule inhibition of STOML3 oligomerization reverses pathological mechanical hypersensitivity. Nat. Neurosci. 20, 209–218 (2017).

26. Haseleu, J., Walcher, J. & Lewin, G. R. The mechanotransduction protein STOML3 is required for proprioceptor plasticity following peripheral nerve regeneration. Exp. Physiol. 10.1113/EP092428 (2025) doi:10.1113/EP092428.

27. Poole, K., Herget, R., Lapatsina, L., Ngo, H. D. & Lewin, G. R. Tuning Piezo ion channels to detect molecular-scale movements relevant for fine touch. Nat Commun 5, 3520 (2014).

28. Zhou, Z. et al. MDFIC2 is a sensory neuron-specific PIEZO channel auxiliary subunit. Proc. Natl. Acad. Sci. U. S. A. 123, e2530071123 (2026).

29. Patkunarajah, A. et al. TMEM87a/Elkin1, a component of a novel mechanoelectrical transduction pathway, modulates melanoma adhesion and migration. Elife 9, (2020).

30. Murthy, S. E. et al. The mechanosensitive ion channel Piezo2 mediates sensitivity to mechanical pain in mice. Sci. Transl. Med. 10, eaat9897 (2018).

31. Qi, Y. et al. Membrane stiffening by STOML3 facilitates mechanosensation in sensory neurons. Nat. Commun. 6, 8512 (2015).

32. Umlauf, E., Mairhofer, M. & Prohaska, R. Characterization of the Stomatin Domain Involved in Homo-oligomerization and Lipid Raft Association. J. Biol. Chem. 281, 23349– 23356 (2006).

33. Brand, J. et al. A stomatin dimer modulates the activity of acid-sensing ion channels. EMBO J. 10.1038/emboj.2012.203 (2012) doi:10.1038/emboj.2012.203.

34. Mannsfeldt, A. G., Carroll, P., Stucky, C. L. & Lewin, G. R. Stomatin, a MEC-2 like protein, is expressed by mammalian sensory neurons. Mol. Cell. Neurosci. 13, 391–404 (1999).

35. Agostinelli, E. et al. A Role for STOML3 in Olfactory Sensory Transduction. eneuro 8, ENEURO.0565-20.2021 (2021).

36. Goldstein, B. J., Kulaga, H. M. & Reed, R. R. Cloning and characterization of SLP3: a novel member of the stomatin family expressed by olfactory receptor neurons. J. Assoc. Res. Otolaryngol. JARO 4, 74–82 (2003).

37. Liang, X., Taylor, M., Napier-Jameson, R., Calovich-Benne, C. & Norris, A. A Conserved Role for Stomatin Domain Genes in Olfactory Behavior. eNeuro 10, ENEURO.0457-22.2023 (2023).

38. Gao, J. et al. Structures of human organellar SPFH protein complexes. Nat. Commun. 16, 10064 (2025).

39. Stoner, J., Li, S. & Fu, Z. Structural basis for membrane microdomain formation by a human Stomatin complex. Nat. Commun. 16, 7439 (2025).

40. Fu, Z. & MacKinnon, R. Structure of the flotillin complex in a native membrane environment. Proc. Natl. Acad. Sci. 121, e2409334121 (2024).

41. Vallese, F., Li, H., Barazzuol, L., Calì, T. & Clarke, O. B. Stomatin encapsulates aquaporin-1 and urea transporter-B in the erythrocyte membrane. Sci. Adv. 12, eaec1721 (2026).

42. Jia, X., Liu, G., Li, H. & Qian, H. Structural insights into the organization of the human Erlin complex. J. Mol. Cell Biol. 18, mjaf060 (2026).

43. Yan, L. et al. Stomatin organizes laterally segregated membrane microdomains to modulate transmembrane transport and signal transduction. Nat. Commun. 10.1038/s41467-026-78250-0 (2026) doi:10.1038/s41467-026-78250-0.

44. Kadurin, I., Huber, S. & Gründer, S. A single conserved proline residue determines the membrane topology of stomatin. Biochem. J. 418, 587–594 (2009).

45. Rungaldier, S. et al. Structure-function analysis of human stomatin: A mutation study. PLOS ONE 12, e0178646 (2017).

46. Bickel, P. E. et al. Flotillin and Epidermal Surface Antigen Define a New Family of Caveolae-associated Integral Membrane Proteins. J. Biol. Chem. 272, 13793–13802 (1997).

47. Schulte, T., Paschke, K. A., Laessing, U., Lottspeich, F. & Stuermer, C. A. O. Reggie-1 and reggie-2, two cell surface proteins expressed by retinal ganglion cells during axon regeneration. Development 124, 577–587 (1997).

48. Bause, E. Structural requirements of *N* -glycosylation of proteins. Studies with proline peptides as conformational probes. Biochem. J. 209, 331–336 (1983).

49. Gavel, Y. & Heijne, G. V. Sequence differences between glycosylated and non-glycosylated Asn-X-Thr/Ser acceptor sites: implications for protein engineering. Protein Eng. Des. Sel. 3, 433–442 (1990).

50. Huber, T. B. et al. Podocin and MEC-2 bind cholesterol to regulate the activity of associated ion channels. Proc. Natl. Acad. Sci. 103, 17079–17086 (2006).

51. Schurek, E.-M. et al. A Disease-causing Mutation Illuminates the Protein Membrane Topology of the Kidney-expressed Prohibitin Homology (PHB) Domain Protein Podocin. J. Biol. Chem. 289, 11262–11271 (2014).

52. Verkest, C. et al. Cluster nanoarchitecture and structural diversity of PIEZO1 at rest and during activation in intact cells. Sci. Adv. 11, eady8052 (2025).

53. Saotome, K. et al. Structure of the mechanically activated ion channel Piezo1. Nature 554, 481–486 (2018).

54. Wang, L. et al. Structure and mechanogating of the mammalian tactile channel PIEZO2. Nature 573, 225–229 (2019).

55. Ridone, P. et al. Disruption of membrane cholesterol organization impairs the activity of PIEZO1 channel clusters. J. Gen. Physiol. 152, e201912515 (2020).

56. Ly, A. T. et al. Single-particle tracking reveals heterogeneous PIEZO1 diffusion. Biophys. J. 124, 4457–4470 (2025).

57. Qiao, Z. et al. Cryo-EM structure of the entire FtsH-HflKC AAA protease complex. Cell Rep. 39, 110890 (2022).

58. Lange, F. et al. In situ architecture of the human prohibitin complex. Nat. Cell Biol. 27, 633– 640 (2025).

59. Cabantous, S. et al. A new protein-protein interaction sensor based on tripartite split-GFP association. Sci Rep 3, 2854 (2013).

60. Hyun, S., Maruri Avidal, L. & Moss, B. Topology of Endoplasmic Reticulum Associated Cellular and Viral Proteins Determined with Split GFP. Traffic 16, 787–795 (2015).

61. Sahl, S. J. et al. Direct optical measurement of intramolecular distances with angstrom precision. Science 386, 180–187 (2024).

62. Gwosch, K. C. et al. MINFLUX nanoscopy delivers 3D multicolor nanometer resolution in cells. Nat. Methods 17, 217–224 (2020).

63. Schaefer, I. et al. Protein kinase A mediates modality-specific modulation of the mechanically-gated ion channel PIEZO2. J. Biol. Chem. 104782 (2023) doi:10.1016/j.jbc.2023.104782.

64. Verkest, C. et al. Intrinsically disordered intracellular domains control key features of the mechanically-gated ion channel PIEZO2. Nat. Commun. 13, 1365 (2022).

65. Ershov, D. et al. TrackMate 7: integrating state-of-the-art segmentation algorithms into tracking pipelines. Nat. Methods 19, 829–832 (2022).

66. Tinevez, J.-Y. et al. TrackMate: An open and extensible platform for single-particle tracking. Methods 115, 80–90 (2017).

67. Wagner, T., Kroll, A., Haramagatti, C. R., Lipinski, H.-G. & Wiemann, M. Classification and Segmentation of Nanoparticle Diffusion Trajectories in Cellular Micro Environments. PLOS ONE 12, e0170165 (2017).

68. Hughes, C. S. et al. Single-pot, solid-phase-enhanced sample preparation for proteomics experiments. Nat. Protoc. 14, 68–85 (2019).

69. Kong, A. T., Leprevost, F. V., Avtonomov, D. M., Mellacheruvu, D. & Nesvizhskii, A. I. MSFragger: ultrafast and comprehensive peptide identification in mass spectrometry–based proteomics. Nat. Methods 14, 513–520 (2017).

70. Da Veiga Leprevost, F., et al. Philosopher: a versatile toolkit for shotgun proteomics data analysis. Nat. Methods 17, 869–870 (2020).

71. Yu, F., Haynes, S. E. & Nesvizhskii, A. I. IonQuant Enables Accurate and Sensitive Label-Free Quantification With FDR-Controlled Match-Between-Runs. Mol. Cell. Proteomics 20, 100077 (2021).

72. Phipson, B., Lee, S., Majewski, I. J., Alexander, W. S. & Smyth, G. K. Robust hyperparameter estimation protects against hypervariable genes and improves power to detect differential expression. Ann. Appl. Stat. 10, 946–963 (2016).

73. Kolberg, L. et al. g:Profiler—interoperable web service for functional enrichment analysis and gene identifier mapping (2023 update). Nucleic Acids Res. 51, W207–W212 (2023).

74. Schindelin, J. et al. Fiji: an open-source platform for biological-image analysis. Nat. Methods 9, 676–682 (2012).

75. Perez-Riverol, Y. et al. The PRIDE database at 20 years: 2025 update. Nucleic Acids Res. 53, D543–D553 (2025).

